# The Aliment to Bodily Condition knowledgebase (ABCkb): A database connecting plants and human health

**DOI:** 10.1101/2021.03.20.436160

**Authors:** Aaron Trautman, Richard Linchangco, Rachel Walstead, Jeremy J Jay, Cory Brouwer

## Abstract

**Objective:** Overconsumption of processed foods has led to an increase in chronic diet-related diseases such obesity and type 2 diabetes. Although diets high in fresh fruits and vegetables are linked with healthier outcomes, the specific mechanisms for these relationships are poorly understood. Experiments examining plant phytochemical production and breeding programs, or separately on the health effects of nutritional supplements have yielded results that are sparse, siloed, and difficult to integrate between the domains of human health and agriculture. To connect plant products to health outcomes through their molecular mechanism an integrated computational resource is necessary.

**Results:** We created the Aliment to Bodily Condition Knowledgebase (ABCkb) to connect plants to human health by creating a stepwise path from plant → plant product → human gene → pathways → indication. ABCkb integrates 11 curated sources as well as relationships mined from Medline abstracts by loading into a graph database which is deployed via a Docker container. This new resource, provided in a queryable container with a user-friendly interface connects plant products with human health outcomes for generating nutritive hypotheses. All scripts used are available on github (https://github.com/atrautm1/ABCkb) along with basic directions for building the knowledgebase.

## Introduction

The growth of obesity worldwide correlates strongly with overconsumption of processed foods [1].This has contributed to an increase in chronic diet-related diseases like type 2 diabetes (T2D), heart disease, and some cancers [2]. Exercise and diets high in fruit, vegetables, whole grains, and nuts have been linked with healthier outcomes and reduce the risk of developing these diseases [3]. Unfortunately, the specific mechanisms driving these associations are poorly understood. The Plant Pathways Elucidation Project (P2EP) was a collaboration started to uncover the mechanisms between plant-pathway products and human health [4]. Three questions drove this collaboration: “What do plants make,” “How do they make them,” and “What is their effect on human health?” The ABCkb was developed to capture the information required to answer these questions and provide researchers with a tool to build informed, nutritive hypotheses with molecular mechanisms as the linking factor between dietary plants and human health.

These questions closely align to the recently released “2020–2030 Strategic Plan for NIH Nutrition Research.”. This plan contains 4 strategic goals for further study to move closer to a precision nutrition approach including foundational research into “What do we eat and how does it affect us?” as well as understanding “How can we improve the use of food as medicine?” A cornerstone for answering these questions and the questions of the P2EP collaboration is an understanding of the mechanism of action of how our diet affects our health.

However, manually capturing this information is a difficult, time-consuming task due to scattered bodies of scientific knowledge. Currently available resources contain partial information to answer these questions, but they do not address mechanism of action. For example, the Comparative Toxicogenomics Database (CTD) connects chemicals to human health through human genes by manually curating associations between chemicals, genes, pathways and phenotypes but excludes nutritional data [5]. Specialized nutritional databases like FooDB (https://foodb.ca) and Phenol-Explorer aid researchers in estimating quantity of phytochemical content, but lack human phenotypic information [6]. NutriChem was developed to bridge the gap between plant-based nutrition and human disease through the chemicals contained in those plants, but does not contain gene-chemical associations, a key part of the driving molecular mechanisms between diet and human health [7]. While a small proportion of assertions are in available databases, others are hidden in published research and can only be extracted through extensive reading or by natural language processing (NLP) the literature. Given the rise in diet-related diseases, and the pursuit of personalized nutrition, an integrated resource to develop nutritive hypotheses is necessary.

## Main Text

We have developed the Aliment to Bodily Condition Knowledgebase (ABCkb) to address the gap of connecting plant compounds to human indications through their mechanism of action. The ABCkb integrates multiple resources for building informed hypotheses with molecular mechanisms as the linking factor between dietary plants and human health. To accomplish this, the ABCkb uses both structured and unstructured data sources (Fig. 1). The structured resources are publicly accessible, curated databases and the unstructured data is in the form of Medline abstracts. Data is extracted, transformed, and then loaded into a Neo4j database. To help users begin discovering easily, the knowledgebase is available as a Docker instance. Additionally, an interface is provided to aid discovery of nutritive connections.

**Figure 1.**
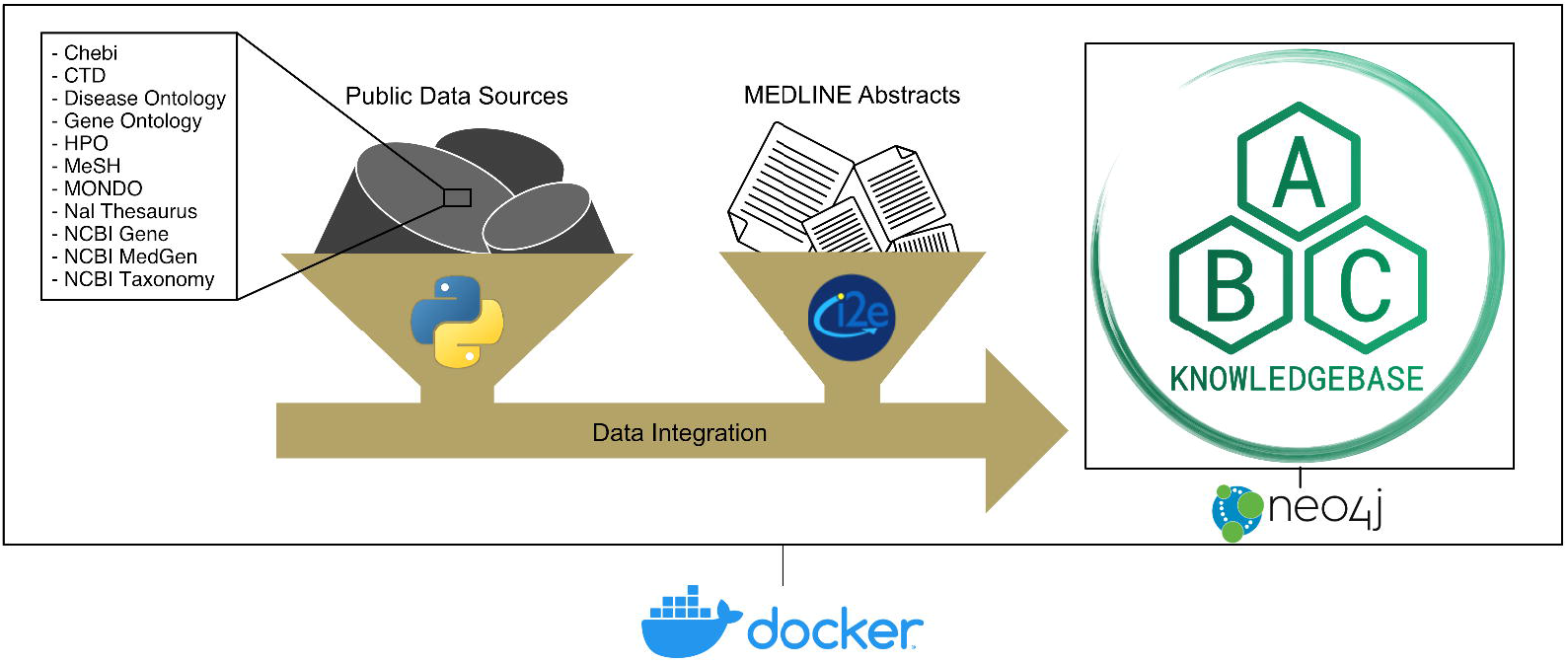
ABCkb Pipeline Overview. The architectural diagram of our Knowledgebase shows the various tools and resources utilized to generate the database.

### Structured resource collection

Structured data from 11 resources (Fig. 2) produce five major labels (Plant, Chemical, Gene, Pathway, Phenotype) in a Neo4j graph database. Connections between these labels are provided by both structured data, and unstructured MEDLINE Abstracts through NLP. The ABCkb utilizes three types of structured data sources: ontologies, structured vocabularies, and databases.

**Figure 2.**
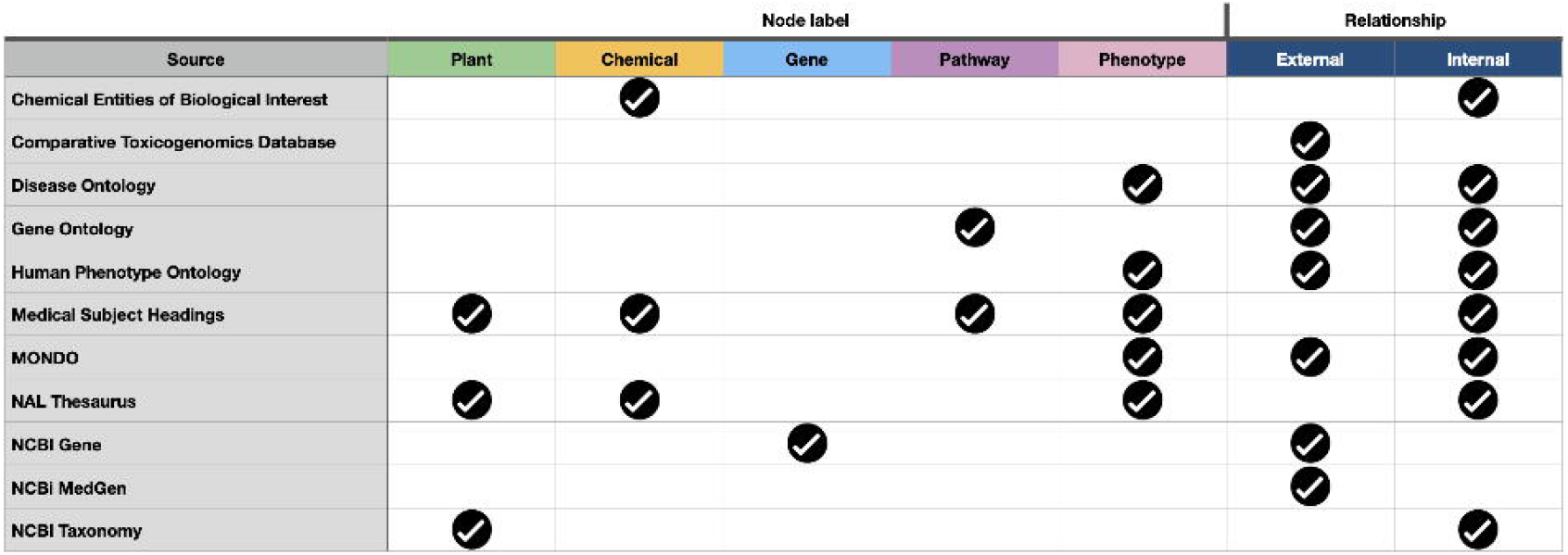
ABCkb data sources. Data from each source is transformed into one of the 5 labels and may provide external and internal references to nodes within the knowledgebase. The CTD provides manually curated references between labels with no original node labels.

#### Ontologies and Structured Vocabularies

The ontologies and structured vocabularies enable well-controlled links between chemicals, pathways, and phenotypes. The Chemical Entities of Biological Interest provide chemical nodes and semantic connections between chemicals [8]. Genes are grouped into pathways from the Gene Ontology resource [9, 10]. Human phenotypes are represented from three sources. The Disease Ontology categorizes human diseases with phenotypic characteristics [11]. The Human Phenotype Ontology provides phenotypic abnormalities not found within the Disease Ontology which allows researchers to focus on specific phenotypic symptoms and the associated molecular mechanisms [12]. Finally, the MONDO Disease Ontology was used to connect phenotype nodes from multiple sources [13]. The Medical Subject Headings resource provided nodes and connections for all major labels with the exception of Genes [14]. Additional plant, chemical, and phenotype nodes are extracted from the National Agricultural Library Thesaurus [15]. Both NAL and MeSH are general sources of knowledge for their domains, provide a wide range of term synonyms, and are used in NLP queries to extract associations from literature and connect extracted associations in the ABCkb. Incorporation of these sources expands the diet to disease connections within the ABCkb

#### Databases

Several databases were utilized to increase molecular mechanisms from plant to human disease in the ABCkb. The Comparative Toxicogenomics database was brought in for over 7.4 million manually curated associations between chemicals, genes, pathways, and phenotypes [5]. The National Center for Biotechnology Information provides various databases available to the public of which, we utilized three. All plants under the Embryophyta clade from the NCBI Taxonomy database produce plant nodes and phylogenetic relationships between plants [16, 17]. The Gene database provides gene names, types, and synonyms [18]. Finally, relationships between NCBI gene nodes and MONDO phenotypes were extracted from the NCBI MedGen database [19]. The compendium of structured data sources provide molecular pathways from diet to disease. However, unstructured literature contains informative relationships not contained within these sources.

### Unstructured NLP collection

To uncover associations in literature, elucidate molecular mechanisms, and answer the three questions of the P2EP, we mined the literature using Linguamatics’ I2E NLP text mining platform. This platform utilizes ontologies and structured vocabularies to transform unstructured text into structured assertions.

#### Natural Language Processing of MEDLINE Abstracts

The I2E platform employs a graphical user interface to aid NLP query development, where each query extracts a set of subjects, objects, and predicates from user-specified ontologies and structured vocabularies. From published abstracts, and titles extracted from MEDLINE in May, 2019, NLP queries were developed with I2E for each of the 4 steps (plant to chemical, chemical to gene, gene to pathway, pathway to phenotype) with an additional query from genes to phenotypes. All I2E assertions generated are provided to users of the ABCkb as source files and are parsed when the graph database is built.

### Statistics and application

Extracted public data sources generated over 957,000 nodes with over 11 million relationships. NLP results from I2E queries make up 1.26 million of the overall relationship count, with 1.25 million novel relationships not from structured public data sources. There are a total of 14 node labels, with the primary focus on 5 labels forming a direct path from plants to human health (Fig. 3a), and 29 relationship types from 11 different sources (Fig. 3b).

**Figure 3.**
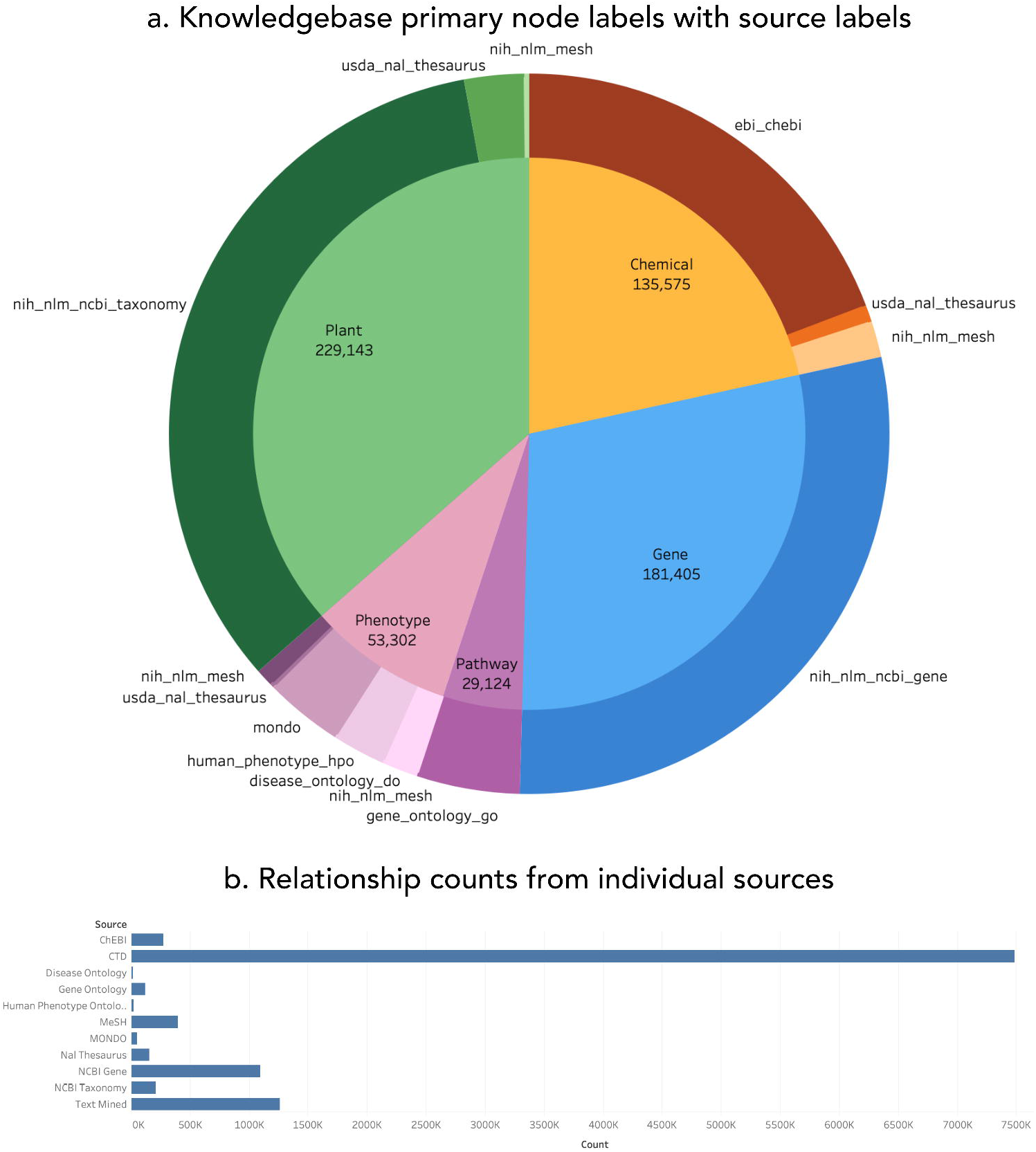
ABCkb node and relationship statistics. a. The pie chart shows primary labels indicated by color with named secondary (source) labels, shaded and sized by proportion of total nodes in the knowledgebase. b. The sum of relationship counts for each source is indicated by the bar chart.

The collection of nodes and relationships in the ABCkb provides novel connections between dietary plants and human phenotypes, and further clarifies connections found in public resources. This is accomplished through the incorporation of molecular associations extracted from literature and structured resources where available. Information in the network flows from plants to phenotypes with assertions in both directions, which allows for query flexibility. Start and end node types are not enforced which allows queries from any point, to any point. To explore the database and discover connections, users have 3 choices: use the provided interface, build graphs with the built in Neo4j interface, or navigate the knowledgebase through a terminal.

The provided user-friendly interface aids users unfamiliar with Neo4j query language (Cypher) to browse the contents within and examine nutritive connections (Fig. 4). On the home page, users are provided a search box to enter in a search term. This scans the nodes in the knowledgebase and returns results ranked by similarity to search term. Users can select nodes and continue to build a query to any end point within the knowledgebase (plant, chemical, gene, pathway, or phenotype). Running the query scans the database for all paths to the selected end point and returns them to the user, which are available to download. Additionally, a Cypher query is available to users that can be used in the built in Neo4j interface or the terminal.

**Figure 4.**
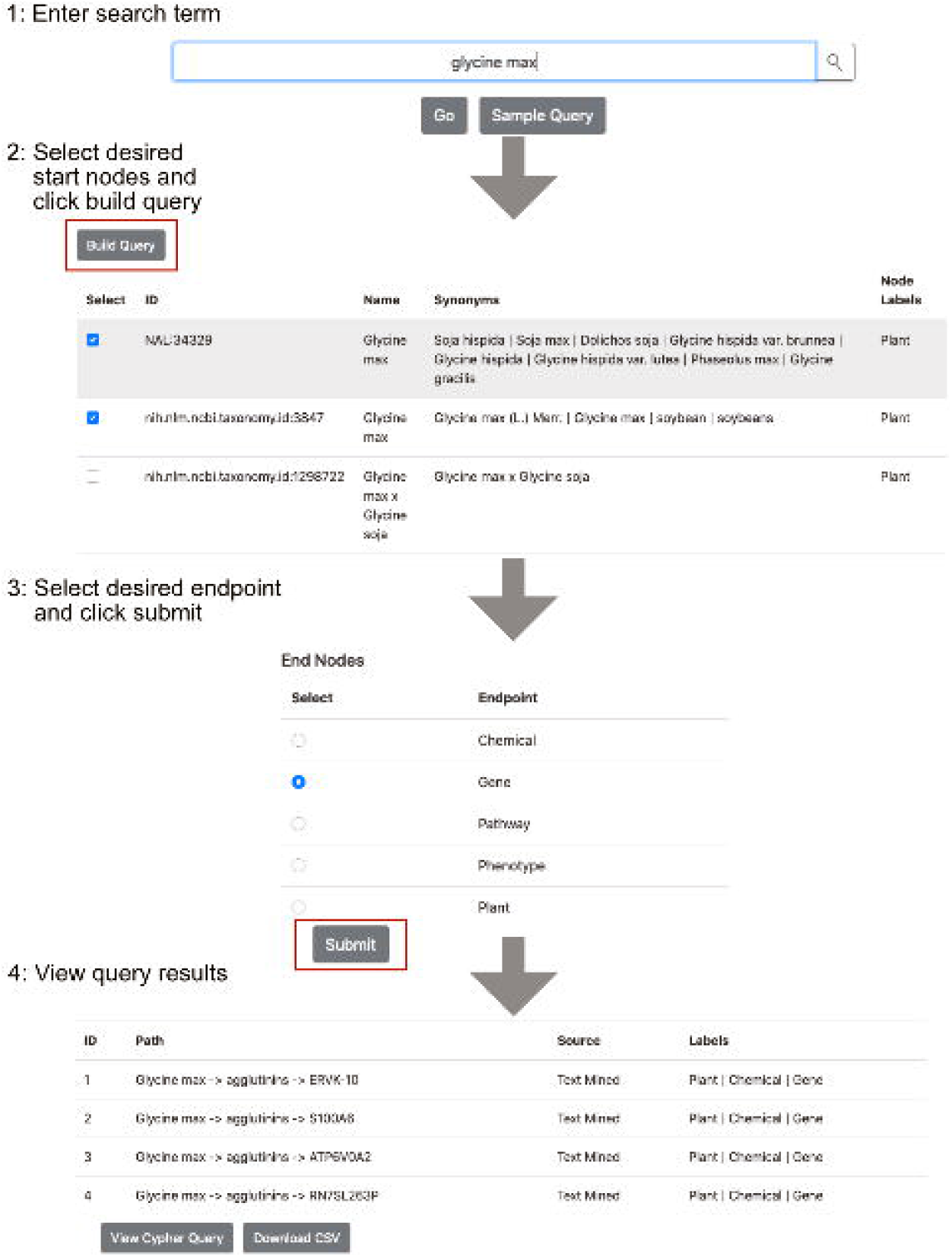
Browsing the ABCkb Interface. There are 4 primary steps to browsing using the provided interface. Once the query endpoint is selected and the user clicks submit, they have the option of downloading all results as a csv, or viewing the Cypher query.

#### Oat and T2D

To demonstrate how the ABCkb connects dietary plants to separate human indications through molecular mechanisms, a graph was created in the ABCkb depicting the diet-disease network between *Avena sativa*, T2D, and heart failure (Fig 5). This network indicates that consumption of oats affects cholesterol levels in the body, which in turn is associated with the gene HSD11B1 that affects lipid metabolic processes with both positive and negative impacts on the incidence of T2D. These relationships are due to the presence of beta-glucan in oat grains. Consumption of beta-glucan-containing oat can help lower LDL cholesterol [20]. The cholesterol lowering effects of oat can also be attributed to the presence of certain lipids and proteins [21]. The proteins in oat with low lysine-arginine and methionine-glycine ratios contribute to lower total cholesterol and LDL cholesterol levels. Hypocholesterolemic properties of oat cannot simply be attributed to one factor, but a combination of many, including oleic acid, vitamin E, and plant sterols [21].

**Figure 5.**
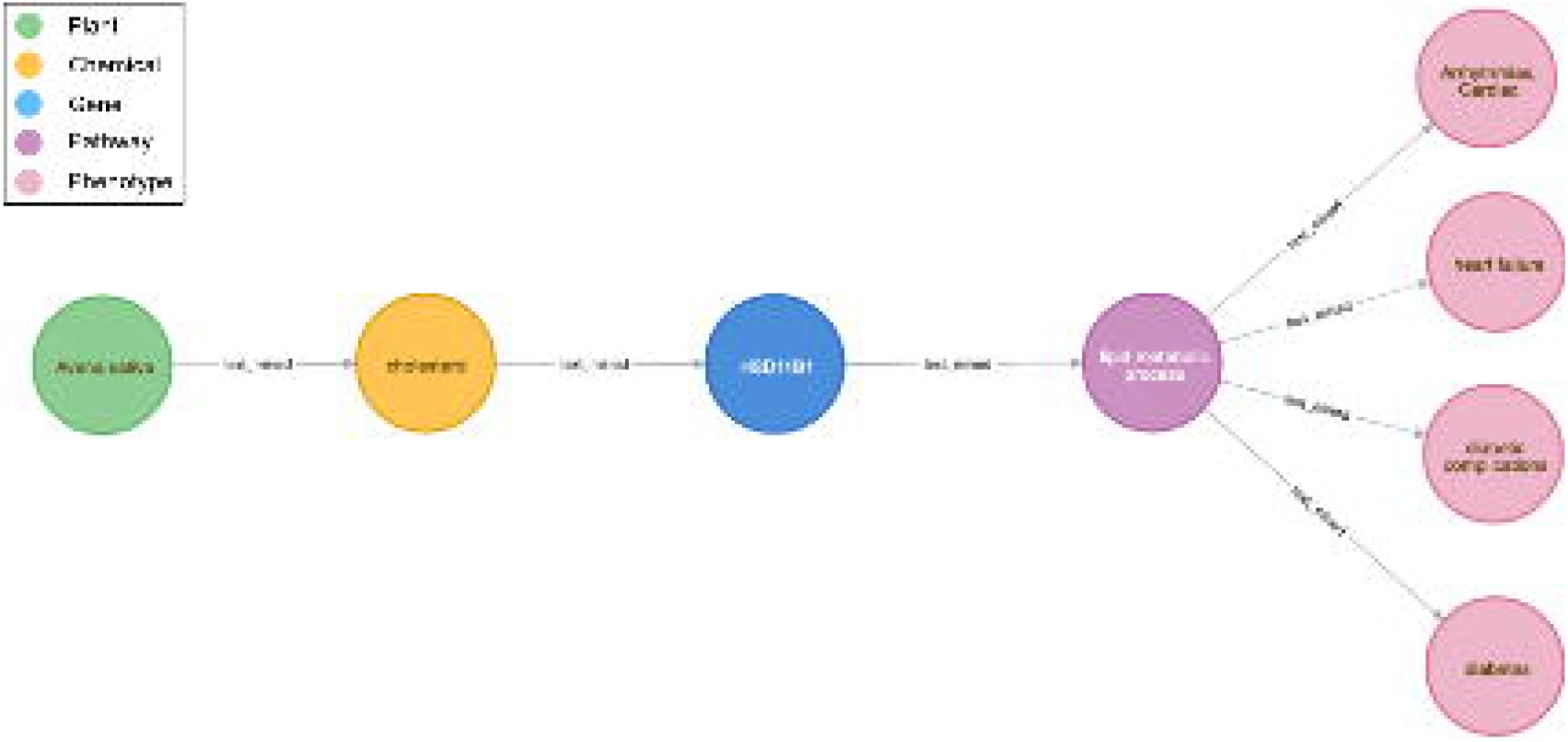
Visualizing the results of Avena sativa to diabetes and heart failure via Hydroxysteroid 11-Beta Dehydrogenase 1. This meta-path highlights the connectivity between oats, diabetes, and heart failure through the gene HSD11B1 from the ABCkb.

T2D patients frequently have abnormal levels of many different lipids, as well as abnormal qualities to these lipids, for example, T2D patients experience normal or slightly elevated LDL cholesterol with increased LDL oxidation and glycation [22]. Dyslipidemia in T2D patients is associated with cardiovascular disease [23, 24]. This creates an elevated risk for cardiovascular diseases including atherosclerosis, and dislipidemia may play a role in these risks [24]. In the graph, HSD11B1 is the human gene connecting this relationship. HSD11B1 expression is increased in adipose tissues of obese individuals [25]. Dysregulation of HSD11B1 is associated with an imbalance of glucocorticoid in adipose tissues, glucose imbalance, and visceral fat accumulation [26]. These factors contribute to metabolic syndrome, which puts patients at a higher risk for cardiac diseases [27]. Various SNPs in HSD11B1 have associations with T2D, metabolic syndrome, and hypertension [28, 29, 30, 31].

Due to the established relationship between oat beta-glucans, cholesterol, and weight, the connection to T2D is logical [20, 26]. Decreased weight, specifically visceral fat in the abdomen, would result in reduced expression of HSD11B1, which would improve regulation of cortisol. Further examination of the oat - cholesterol –HSD11B1 relationship could be very informative to both patients and doctors in making more informed dietary choices and reducing the risk of developing T2D. This example demonstrates the ABCkb ability to connect seemingly separate conditions through the molecular mechanistic links within.

### Discussion

The ABCkb integrates structured and unstructured resources in a network that connects plants to human disease through molecular mechanisms. This reduces the time required to manually connect these links through each individual resource. Additionally, knowledge discovery is aided by the development of a user-friendly interface. All of these components provide precision nutrition a path to better understand the mechanisms behind diet-related conditions. The ABCkb is available from github (https://github.com/atrautm1/ABCkb).

## Limitations

- Microbiota contributions to diet and human disease. Bacteria within the gut are known to affect disease both through the production of metabolites and the conversion of plant phytochemicals. In addition, gut bacteria are affected by diet. Future implementations of the ABCkb will contain microbiota associations to enhance precision nutrition hypotheses.
- Mining abstracts versus full text. Abstracts contain valuable associations, however associations full text articles would provide a greater number of associations.
- Incorporating genomic data. Precision nutrition hypotheses and treatment plans will depend on patient genomic data, to provide optimal dietary solutions for each individual. Future versions of the ABCkb should incorporate human genomic data.

## Ethics approval and consent to participate

Not applicable

## Consent for publication

Not applicable

## Availability of data and materials

All source code is available on the project github (https://github.com/atrautm1/ABCkb) along with basic instructions for building and running the knowledgebase.

The prebuilt neo4j database is available from: 10.6084/m9.figshare.13653539

Operating system(s): Platform independent

Database version: Neo4j 3.5

License: GNU GPL v3

## Abbreviations

ABCkb: Alimentary to Bodily Condition knowledgebase
P2EP: Plant Pathways Elucidation Project
CTD: Comparative Toxicogenomics Database
NLP: Natural Language Processing
LDL: Low-Density Lipoprotein
T2D: Type 2 Diabetes

## Author’s contributions

AT and RL designed and built the ABCkb, AT, RW, and CB wrote the manuscript, JJ and CB provided guidance, all authors provided corrections and feedback. All authors have read and approved the final manuscript.

## Acknowledgements

Steven Blanchard, P2EP program, and the many interns that aided the development of this resource

